# Meta population modelling of narwhals in East Canada and West Greenland - 2017

**DOI:** 10.1101/059691

**Authors:** Lars Witting

## Abstract

This paper combines a catch allocation model for narwhals in East Canada and West Greenland with Bayesian population modelling of the eight summer aggregations of narwhals in the region. The catch allocation model allocates the catches in different hunting areas and seasons to the different summer aggregations, and the population models analyse the impact of these catches on the population dynamics of the eight narwhal aggregations.

The population models run from 1970, and the catch allocation model needs population trajectories from 1970 to the present in order to estimate the catches taken from the different summer aggregations during this period. In an initial run it uses linear transitions between the available abundance estimates; but more elaborate population trajectories are estimated by the fit of the population models to the abundance data. The two models are therefore run in an iterative manner until the catch histories that are estimated by the allocation model, and the abundance trajectories that are estimated by the population models, converge between runs.

Given a converged model and potential future catch options for the different hunts, the model estimates the probabilities of fulfilling management options for the eight summer aggregations of narwhals.

## INTRODUCTION

This paper develops a meta population dynamic model for eight summer aggregations of narwhals in East Canada and West Greenland. It combines a catch allocation model that was developed at the 2014 meeting of the JWG (JWG 2015) with eight population dynamic models, in the aim of assessing the population dynamic implications of hunting.

One of the main challenges in this assessment is the divergence between the population structure of narwhals and the geographical and yearly structure of the hunt. Narwhals return to specific summering grounds and their population structure is best described by the geographical distinctiveness of the different summering grounds; a view that has identified eight populations of narwhal in the Baffin Bay region in East Canada and West Greenland (JWG 2015).

These populations, however, are hunted not only on the summering grounds, but also on their spring and fall migrations in other areas, as well as on the wintering grounds; and these latter areas are typically shared to some degree by individuals from different summer ground.

This lead to the development of a catch allocation model that uses satellite tracking, expert judgment, and abundance estimates to allocate the catches from 24 seasonal and geographically separated hunts between the eight populations of narwhals (JWG 2015). I use this model to produce historical catch histories for the eight summer aggregations, and this allows me to construct population dynamic models that assess the implications of the hunt on the eight summer aggregations. The catch allocation and population dynamics models are then integrated to provide a tool that can assess the sustainability of potential future catches in the 24 hunts, as defined by the impacts of the hunts on the eight summer aggregations of narwhals.

## CATCH ALLOCATION MODEL

The catch allocation model was developed by JWG (2015), and it allocates catches taken in different hunting regions and seasons to the eight summer aggregations of narwhals in East Canada and West Greenland.

The model uses a proportional availability matrix (Table 2 & 1) to describe the availability of the narwhals from the different summer aggregations to the hunts in the different regions and seasons, and it uses a catch matrix (Tables 10 and 11) to describe the annual total removals (catches plus loss) in the different hunts.

**Table 2:**
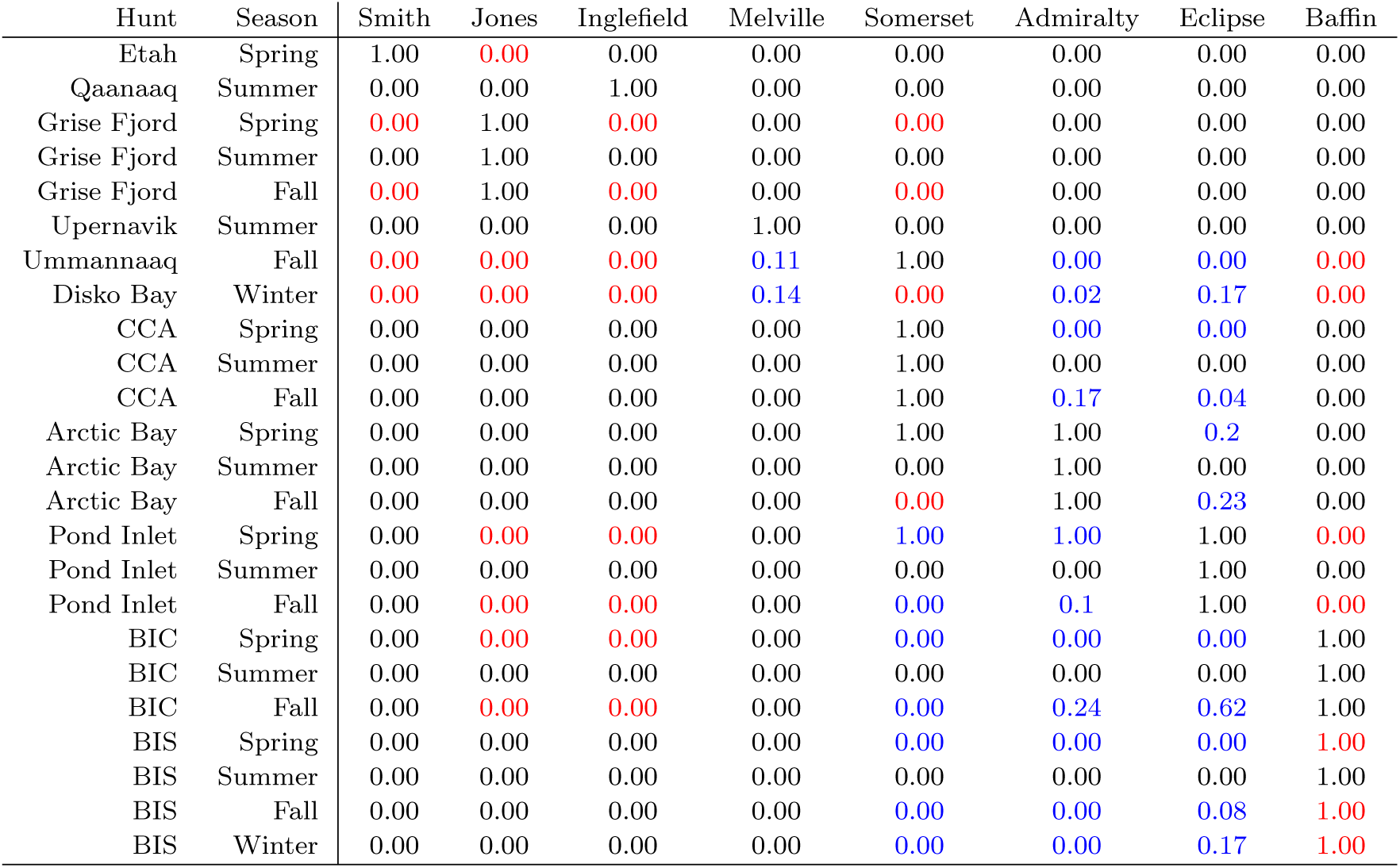
The proportional availability matrix (**P**) of narwhals from summer aggregations to hunting regions. Black numbers are fixed, blue and red are point estimates of beta distributions; red for sensitivity only.

**Table 1:** The proportional availability matrix (**P**) of narwhals from summer aggregations to hunting regions [*x*/*y*: available (*x*) over total (*y*)]. Black numbers (defined zeros and hunts) are fixed, blue (partial hunts) and red (probable zeros and hunts) are beta(α, *β*) distributions (α = *x*+1; *β* = y − *x* + 1); red for sensitivity by changes in *n*.

| Hunt | Season | Smith | Jones | Inglefield | Melville | Somerset | Admiralty | Eclipse | Baffin |
| --- | --- | --- | --- | --- | --- | --- | --- | --- | --- |
| Etah | Spring | 1 | 0/*n* | 0 | 0 | 0 | 0 | 0 | 0 |
| Qaanaaq | Summer | 0 | 0 | 1 | 0 | 0 | 0 | 0 | 0 |
| Grise Fjord | Spring | 0/*n* | 1 | 0/*n* | 0 | 0/*n* | 0 | 0 | 0 |
| Grise Fjord | Summer | 0 | 1 | 0 | 0 | 0 | 0 | 0 | 0 |
| Grise Fjord | Fall | 0/*n* | 1 | 0/*n* | 0 | 0/*n* | 0 | 0 | 0 |
| Upernavik | Summer | 0 | 0 | 0 | 1 | 0 | 0 | 0 | 0 |
| Ummannaq | Fall | 0/*n* | 0/*n* | 0/*n* | 1/9 | 1 | 0/42 | 0/26 | 0/*n* |
| Disko Bay | Winter | 0/*n* | 0/*n* | 0/*n* | 1/7 | 0/*n* | 1/42 | 1/6 | 0/*n* |
| CCA | Spring | 0 | 0 | 0 | 0 | 1 | 0/4 | 0/5 | 0 |
| CCA | Summer | 0 | 0 | 0 | 0 | 1 | 0 | 0 | 0 |
| CCA | Fall | 0 | 0 | 0 | 0 | 1 | 7/42 | 1/26 | 0 |
| Arctic Bay | Spring | 0 | 0 | 0 | 0 | 1 | 1 | 1/5 | 0 |
| Arctic Bay | Summer | 0 | 0 | 0 | 0 | 0 | 1 | 0 | 0 |
| Arctic Bay | Fall | 0 | 0 | 0 | 0 | 0/*n* | 1 | 6/26 | 0 |
| Pond Inlet | Spring | 0 | 0/*n* | 0/*n* | 0 | 2/2 | 4/4 | 1 | 0/*n* |
| Pond Inlet | Summer | 0 | 0 | 0 | 0 | 0 | 0 | 1 | 0 |
| Pond Inlet | Fall | 0 | 0/*n* | 0/*n* | 0 | 0/14 | 4/42 | 1 | 0/*n* |
| BIC | Spring | 0 | 0/*n* | 0/*n* | 0 | 0/2 | 0/4 | 0/6 | 1 |
| BIC | Summer | 0 | 0 | 0 | 0 | 0 | 0 | 0 | 1 |
| BIC | Fall | 0 | 0/*n* | 0/*n* | 0 | 0/5 | 10/42 | 16/26 | 1 |
| BIS | Spring | 0 | 0 | 0 | 0 | 0/2 | 0/4 | 0/6 | *n*/*n* |
| BIS | Summer | 0 | 0 | 0 | 0 | 0 | 0 | 0 | 1 |
| BIS | Fall | 0 | 0 | 0 | 0 | 0/5 | 0/42 | 2/26 | *n*/*n* |
| BIS | Winter | 0 | 0 | 0 | 0 | 0/2 | 0/42 | 1/6 | *n*/*n* |

To allocate the catches from the different hunts to the different summer aggregations, the model needs an additional matrix that describes the abundance in the different stocks per year. These abundance estimates are needed to estimate the relative availability of the different stocks to the different hunts, so that the catches can be allocated to the summer aggregations.

### Allocation matrix

The allocation model was developed in the form of a 24 rows by 8 columns allocation matrix. The eight columns are the individual summer aggregations, and the 24 rows represent hunts divided by 10 regions with some of the hunts divided by season. For each summer aggregation and hunt there is a cell in the matrix, and the matrix is devised so that when multiplied by a vector of removals, the resulting vector will determine the total removals from each summer aggregation.

Each cell of the allocation matrix, **A**, has a value

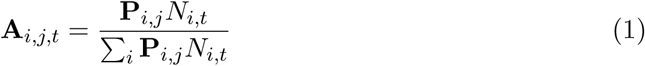
where, **A***_i,j,t_* is the proportion of the jth hunt that is assigned to the *i*th summer aggregation in year *t*, **P***_i,j_* is the proportional availability of the ith summer aggregation to the *j*th hunt, and *N_i,t_* is the abundance of the *i*th summer aggregation in year *t*.

This model assumes that for each summer aggregation there is a proportion between zero and one, **P***_i,j,_* that is available to hunters during the hunting period on the hunting grounds. Each individual that is available is then at equal risk of being taken in the hunt. The sum of the **A***_i,j,t_* should be 1 for each row of **A***_t_*.

To set up the proportional availability matrix (**P**; Table 2 & 1) the JWG reviewed each cell so that each cell in the matrix was given one of five designations:

**Defined zero:** This designated cells that represented improbable situations such as a summer harvest that was not at a summering ground (e.g. a narwhal harvested in summer in Resolute could not come from the Smith Sound summer aggregation and would be assigned to Somerset Island stock) and to hunts in areas that could not have originated in a particular summering ground based on known movements.
**Probable zero:** This designated cells in which a summering aggregation was unlikely to be hunted but proximity during the hunting season, or a presumed migration route did not rule out possible catches and designated cells with no tag data.
**Partial hunt:** This designated cells with tag data showing a portion of the summering aggregation was available or not to hunters.
**Probable hunt:** This designated cells in which a summering aggregation was likely to be fully available to a hunt, based on its geographical proximity to a summering ground or migration route, but for which there was no quantitative evidence such as tag data.
**Defined hunt:** This designates cells representing hunts on summering grounds or known wintering areas of stocks.

Each cell in **P** was then parameterised as a beta(*α, β*) distribution, where *α*= *x* + 1 and *β* = *y* − *x* + 1 would depend on the designation of the hunt. Defined zeros where obtained by *x* = 0 and *y* = 9999, defined hunts by *x* = 9999 and *y* = 9999, probable zeros by *x* = 0 and *y* = *n*, probable hunts by *x* = *y* = *n*, and for a partial hunt that was parameterised by the seasonal and geographical distribution of *y* sattelited trackes individuals, *x* would be the number of the individuals that migrated to the hunting ground. The simulations in this paper, however, set *n* = 9999 which implies that they do not distinguish between probable and defined designations.

### The generation of catch histories

The abundance matrix in the initial run of the model is constructed as linear transitions between the abundance estimates in the abundance estimate matrix of Table 3. In subsequent runs, the abundance matrix is given by the abundance trajectories that the previous runs of the population dynamic models are estimating for the different summer aggregations of narwhals, given the catch histories that were estimated by the previous run of the allocation model. This iterative running of the two models was then conducted three to five times to ensure convergence of the catch histories and abundance trajectories.

**Table 3:** Abundance estimates with CVs for summer aggregations of narwhal.

| Year | Smith | Jones | Inglefield | Melville | Somerset | Admiralty | Eclipse | Baffin |
| --- | --- | --- | --- | --- | --- | --- | --- | --- |
| 1975 | - | - | - | - | - | 28260; 0.22 | - | - |
| 1981 | - | - | - | - | 32520; 0.1 | - | - | - |
| 1985 | - | - | - | - | - | 16400; 0.43 | - | - |
| 1986 | - | - | 8710; 0.25 | - | - | - | - | - |
| 1996 | - | - | - | - | 45360; 0.35 | - | - | - |
| 2001 | - | - | - | - | - | - | - | - |
| 2002 | - | - | - | - | 35810; 0.43 | - | - | - |
| 2003 | - | - | - | - | - | 5360; 0.5 | - | 10070; 0.31 |
| 2004 | - | - | - | - | - | - | 20230; 0.36 | - |
| 2007 | - | - | 8370; 0.25 | 6020; 0.86 | - | - | - | - |
| 2009 | - | - | - | - | - | - | - | - |
| 2010 | - | - | - | - | - | 18050; 0.22 | - | - |
| 2012 | - | - | - | 2980; 0.39 | - | - | - | - |
| 2013 | 16360; 0.65 | 12690; 0.33 | - | - | 49770; 0.2 | 35040; 0.42 | 10490; 0.24 | 17560; 0.35 |
| 2014 | - | - | - | 3090; 0.5 | - | - | - | - |

In either case, for a given year *t*, we would have abundance estimates with cv’s for all summer aggregations. Distributions of catch histories for the summer aggregations during these iterations were generated by a large number of runs of the allocation model over the entire catch history with yearly random draws of each abundance estimate (lognormal based on point estimate and cv) and a **P**_*t*_ matrix drawn from the underlying beta distributions.

## POPULATION DYNAMIC MODELS

Separate population models with density regulated growth were constructed for each of the eight summer aggregations of narwhals. All the models were based on an age and sex structured Bayesian modelling framework that have been used in earlier applications for walrus (Witting and Born 2005, 2013), large cetaceans (Witting 2013), beluga (Heide-Jørgensen et al. In press), and narwhal (e.g., Witting and Heide-Jørgensen 2012a,b,c).

Some of the summering aggregations have only been surveyed once, and a full age-structured model is clearly over parameterized for these cases if the main purpose was parameter estimation by maximum likelihood. Yet, the main purpose here is instead to use a Bayesian framework to integrate prior knowledge on the life history biology of narwhals with survey estimates of abundance for the construction of realistic population dynamic models.

Witting (2009) used the case of beluga in West Greenland to analyse for influence of model uncertainty in the construction of realistic population models in Bayesian assessments of density regulated growth. Assessments were made for one age-structured and four structurally different discrete models, with all assessments being based on the same data. All models gave very similar estimates of current abundance and current production levels, making the choice of model basically a matter of taste.

I have chosen an age and sex structured framework that allows the model to be constructed directly from our prior knowledge of the life history of narwhals. This allows also for a later inclusion of sex structured catches, and age-structured catch data, as it has been done in earlier assessments of Greenland narwhals (Witting and Heide-Jørgensen 2012a,b,c). Age-structured data, however, were not included here, as I wanted to keep the population dynamic models relatively simple (and fast to simulate) in this first run of the meta model.

Let *x* = 15 be the maximum lumped age-class of the model. Let the number 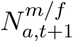 of males (*m*) and females (*f*) in age-classes 0 < *a* < *x* in year *t* + 1 be

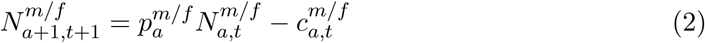
and the number of animals in age-class *x* be

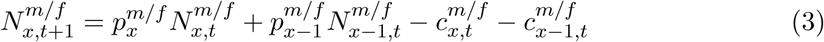
where 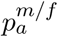 is the age specific survival rate of males/females, and 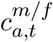 is the age specific catch of males/females in year *t*. The age and gender (*g*) dependent survival rates 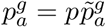 are given as a product between a survival scalar *p* and a relative 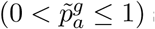 survival rate, with relative survival being one for males and females older than one year of age. The age and gender specific catches 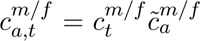 in year *t* is given as a product between the total catch of males/females 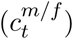, as specified by the catch history, and an age-specific catch selectivity 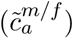 that is uniform except that no animals are taken from age-class zero.

The number of females and males in age-class zero is 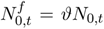 and 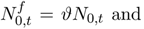, where *ϑ* is the fraction of females at birth, and

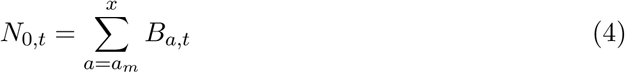
where *a_m_* is the age of the first reproductive event and *B_a,t,_* the number of births from females in age class *a*, is

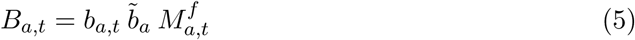
where *b_a,t_* is the birth rate in year *t* for age-class *a* females should they be at their age-specific reproductive peak, 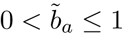 is the relative age-specific birth rate (1 for all mature females), and 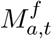 is the number of mature females in age-class *a* in year *t*, defined as

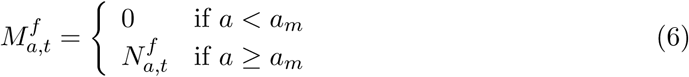
Let *b_a,t_* be

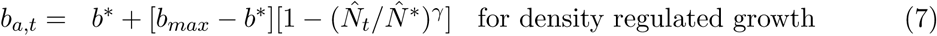
where *b*\* is the birth rate at population dynamic equilibrium (assuming zero catch and equilibrium denoted by *), *b_max_* is the maximal birth rate, γ is the density dependence parameter, and the abundance component that imposes density dependence is the one-plus component

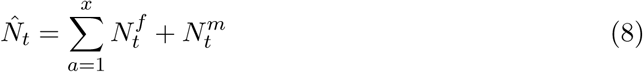
Given a stable age-structure and no catch, let, for a traditional model of exponential or density regulated growth, λ be a constant defined by 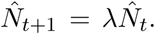. The sustainable yield is then 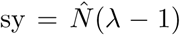, and for the density regulated model there is an optimum 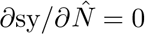; the maximum sustainable yield (msy) at 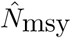, also known by the maximum sustainable yield rate (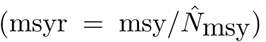) and the maximum sustainable yield level (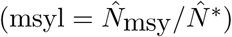).

## STATISTICAL METHODS

The assessment model was fitted to data by projecting the population under the influence of the historical catches. A Bayesian statistical method (e.g, Berger 1985; Press 1989) was used, and posterior estimates of model parameters and other management related outputs were calculated. This implied an integration of the product between a prior distribution for each parameter and a likelihood function that links the probability of the data to the different parameterisations of the model.

### Prior distributions

All models had the same priors on the biological parameters (see Table 4), and they were all initiated in 1970. All the summer aggregations with only one or two abundance estimates available (Smith, Jones, Eclipse, Baffin) and Admiralty seems to have had a very low exploitation rate in the beginning of the period, so for these I assumed that the population was close to the carrying capacity in 1970. For the remaining aggregations (Inglefield, Melville and Somerset), with a somewhat larger early exploitation, I assumed that the abundance in 1970 was lower than the carrying capacity.

**Table 4:** Prior distributions for the different models (*M*). The list of parameters: *N*_0_ is the initial abundance, *N*\* the population dynamic equilibrium abundance, *p* the yearly survival, *p*_0_ the first year survival, *b* the birth rate, *a_m_* the age of the first reproductive event, *ϑ* the female fraction at birth, γ the density regulation, *c_h_* the catch history, and *β_i_* the abundance estimate bias (*i*: data reference). Abundance is given in thousands. The prior probability distribution is given by superscripts; *p*: fixed value, *u*: uniform (min,max), *U*: log uniform (min,max), *b*: beta 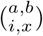 with *i*=min and *x*=*max*, and *f*: file distribution.

| $M$ | $N_0$ | $N^*$ | $p$ | $p_0$ | $b$ | $a_m$ | $\vartheta$ | $\gamma$ | $c_h$ | $\beta_a$ |
| --- | --- | --- | --- | --- | --- | --- | --- | --- | --- | --- |
| smith | - | $2,80^U$ | $\begin{smallmatrix} 2,2 \\ .95,1 \end{smallmatrix}^b$ | $.5,1^u$ | .33 | $\begin{smallmatrix} 2,2 \\ 7,15 \end{smallmatrix}^b$ | .5 | $2,4^u$ | f | - |
| jones | - | $2,60^U$ | $\begin{smallmatrix} 2,2 \\ .95,1 \end{smallmatrix}^b$ | $.5,1^u$ | .33 | $\begin{smallmatrix} 2,2 \\ 7,15 \end{smallmatrix}^b$ | .5 | $2,4^u$ | f | - |
| ingle | $1,25^U$ | $3,30^U$ | $\begin{smallmatrix} 2,2 \\ .95,1 \end{smallmatrix}^b$ | $.5,1^u$ | .33 | $\begin{smallmatrix} 2,2 \\ 7,15 \end{smallmatrix}^b$ | .5 | $2,4^u$ | f | $.01,1^U$ |
| melvi | $.8,20^U$ | $3,30^U$ | $\begin{smallmatrix} 2,2 \\ .95,1 \end{smallmatrix}^b$ | $.5,1^u$ | .33 | $\begin{smallmatrix} 2,2 \\ 7,15 \end{smallmatrix}^b$ | .5 | $2,4^u$ | f | - |
| somer | $5,60^U$ | $25,90^U$ | $\begin{smallmatrix} 2,2 \\ .95,1 \end{smallmatrix}^b$ | $.5,1^u$ | .33 | $\begin{smallmatrix} 2,2 \\ 7,15 \end{smallmatrix}^b$ | .5 | $2,4^u$ | f | - |
| admir | - | $10,40^U$ | $\begin{smallmatrix} 2,2 \\ .95,1 \end{smallmatrix}^b$ | $.5,1^u$ | .33 | $\begin{smallmatrix} 2,2 \\ 7,15 \end{smallmatrix}^b$ | .5 | $2,4^u$ | f | - |
| eclip | - | $5,50^U$ | $\begin{smallmatrix} 2,2 \\ .95,1 \end{smallmatrix}^b$ | $.5,1^u$ | .33 | $\begin{smallmatrix} 2,2 \\ 7,15 \end{smallmatrix}^b$ | .5 | $2,4^u$ | f | - |
| baffi | - | $3,60^U$ | $\begin{smallmatrix} 2,2 \\ .95,1 \end{smallmatrix}^b$ | $.5,1^u$ | .33 | $\begin{smallmatrix} 2,2 \\ 7,15 \end{smallmatrix}^b$ | .5 | $2,4^u$ | f | - |

The catch histories in a run of the population models were estimated by the allocation model over the complete catch history starting in 1970. These distributions were then used as catch priors in the population models, with specific catch histories being drawn from the prior for each iteration of a population model.

These catch priors were constructed from the distribution of possible total removals that was estimated for 2011 by 200 random draws of the allocation model (Figure 1), together with two complete catch histories, a minimum catch history (*c_min_*, represented by the 1th percentile of this distribution over time) and a maximum catch history (*c_max_*, represented by the 99th percentile). These percentiles were estimated from 1000 random draws of the allocation model over the entire catch history. The 2011 distribution was then rescaled to run from zero to one, with a value (*x*) drawn at random from the distribution for each parameterisation of a population model, with the catch history calculated as 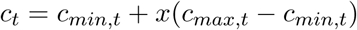

**Figure 1:**
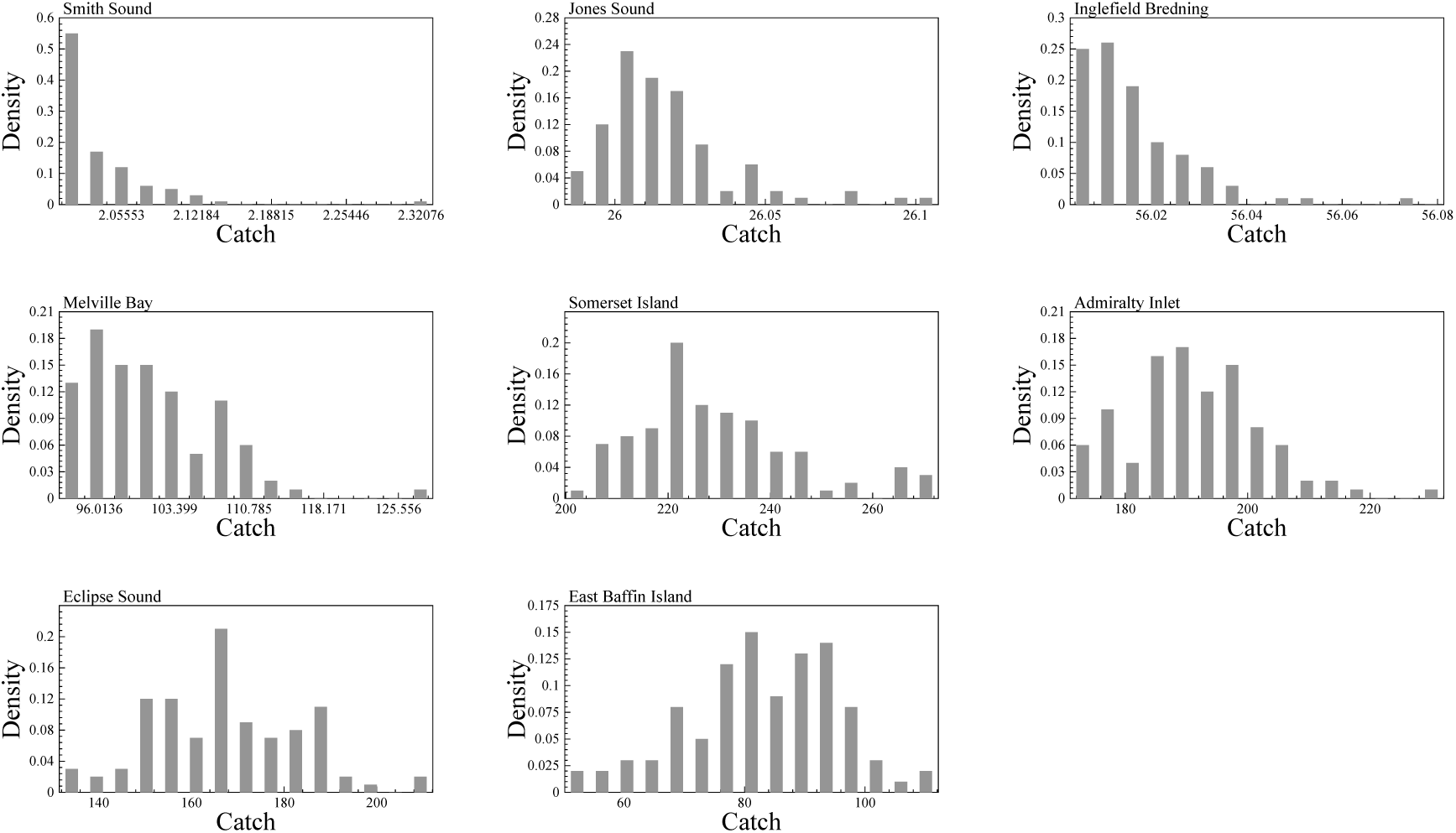
Catch distributions per summer aggregation, estimates for year 2011.

### Bayesian integration

The Bayesian integration was obtained by the sampling-importance-resampling routine (Jeffreys 1961; Berger 1985; Rubin 1988), where *n_s_* random parameterisations *θ_i_* (1 ≤ *i* ≤ *n*_1_) are sampled from an importance function *h*(*θ*). This function is a probability distribution function from which a large number, n_s_, of independent and identically distributed draws of *θ* can be taken. *h*(*θ*) shall generally be as close as possible to the posterior, however, the tails of *h*(*θ*) must be no thinner (less dense) than the tails of the posterior (Oh and Berger 1992). For each drawn parameter set *θ_i_* the population was projected from the first year with a harvest estimate to the present. For each draw an importance weight, or ratio, was then calculated

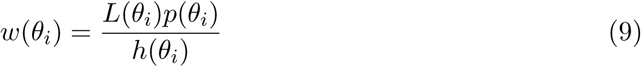
where *L*(*θ_i_*) is the likelihood given the data, and *h*(*θ_i_*) and *p*(*θ_i_*) are the importance and prior functions evaluated at *θ_i_*. In the present study the importance function is set to the joint prior, so that the importance weight is given simply by the likelihood. The *n_s_* parameter sets were then re-sampled *n_r_* times with replacement, with the sampling probability of the *i*th parameter set being

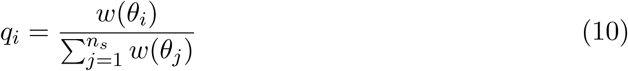

This generates a random sample of the posterior distribution of size *n_r_*.

The method of de la Mare (1986) was used to calculate the likelihood L under the assumption that observation errors are log-normally distributed (Buckland 1992)

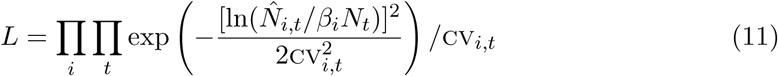
Where 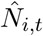 is the point estimate of the ith set of abundance data in year t, cvg_t_ is the coefficient of variation of the estimate, *N_t_* is the simulated abundance, and *β_i_* a bias term with is set to one for absolute abundance estimates.

### Probabilities of meeting objectives

In order to assess the sustainability of future catches on the eight populations of narwhals, I used the Bayesian framework to estimate the probabilities that an assumed management objective would be fulfilled for potential future catches from each population.

Given future annual catches c in the period 2015 to 2020, I applied the objective

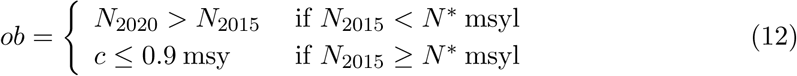
Given the population dynamic model and the data, the probability of meeting the objective *ob* is straightforwardly calculated from the Bayesian statistical method. For each parameterisation *θ_i_* of the random sample of the posterior distribution of size *n*_2_, we have perfect knowledge of the status of the population so that it can be determined if Eq. (12) is true or false. Hence, the probability *p*(*ob*) of meeting the objective is

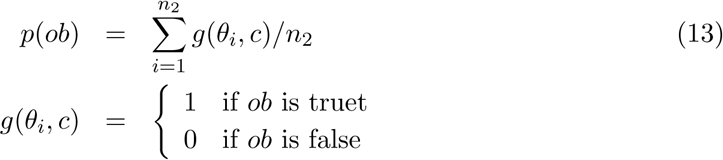
with the sum given over the complete random sample of the posterior distribution.

While the sustainability of the hunt has to be identified at the population level, recommendations on the sustainability of potential future hunts should preferably be addressed in relation to hunting grounds. To achieve this, for a given a set of potential future catches for each hunt, I used the allocation model to calculate the distributions of future catches for the different populations, with these distributions reflecting the uncertainty in the allocation of catches between the populations. Then, by having these distributions, I could for the catches of each percentile of these distributions calculate the probability that the assumed management objective would be fulfilled for the different populations. This allows the set of potential catches for the different hunts to be adjusted until the probabilities of fulfilling management objectives would be above agreed threshold levels for all populations.

## RESULTS

The convergence of the catch and abundance trajectories over the different iterations of the allocation and population dynamic models is shown in Figure 2.

**Figure 2:**
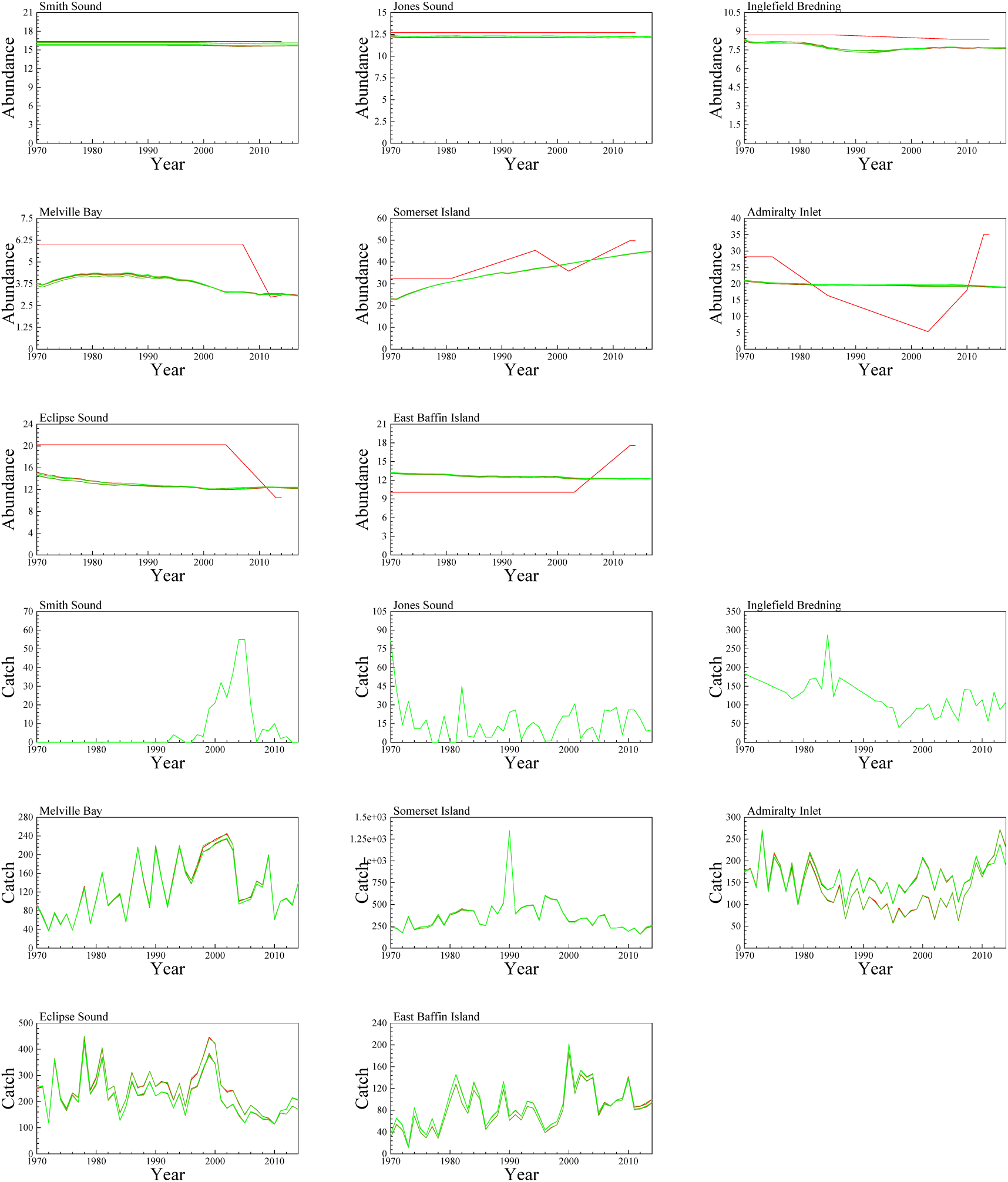
The convergence of the abundance trajectories and catch histories as a function of the number of iterations of the complete meta aggregation model, with iteration number increasing with colour transitions from clear red to clear green. Abundance is given in thousands.

The sampling statistics of the last run of the Bayesian population models are shown in Table 5. The estimated trajectories of the eight summer aggregations are shown in Figure 3, and the posterior parameter estimates in Table 6, with plots of the posterior and realised prior distributions given in Figures 5 to 12. The final estimates of the catch histories per summer aggregation are shown in Figure 4.

**Table 5:** Sampling statistics for the different models (*M*). The number of parameter sets in the sample (*n_S_*) and the resample (*n_R_*), the number of unique parameter sets in the resample, and the maximum number of occurrences of a unique parameter set in the resample. n_S_ and n_R_ are given in thousands.

| $M$ | $n_S$ | $n_R$ | Unique | Max |
| --- | --- | --- | --- | --- |
| smith | 500 | 5 | 4964 | 2 |
| jones | 500 | 5 | 4926 | 2 |
| ingle | 2000 | 5 | 3672 | 11 |
| melvi | 500 | 5 | 4794 | 3 |
| somer | 500 | 5 | 4747 | 4 |
| admir | 500 | 5 | 4915 | 3 |
| eclip | 500 | 5 | 4907 | 2 |
| baffi | 500 | 5 | 4899 | 3 |

**Figure 3:**
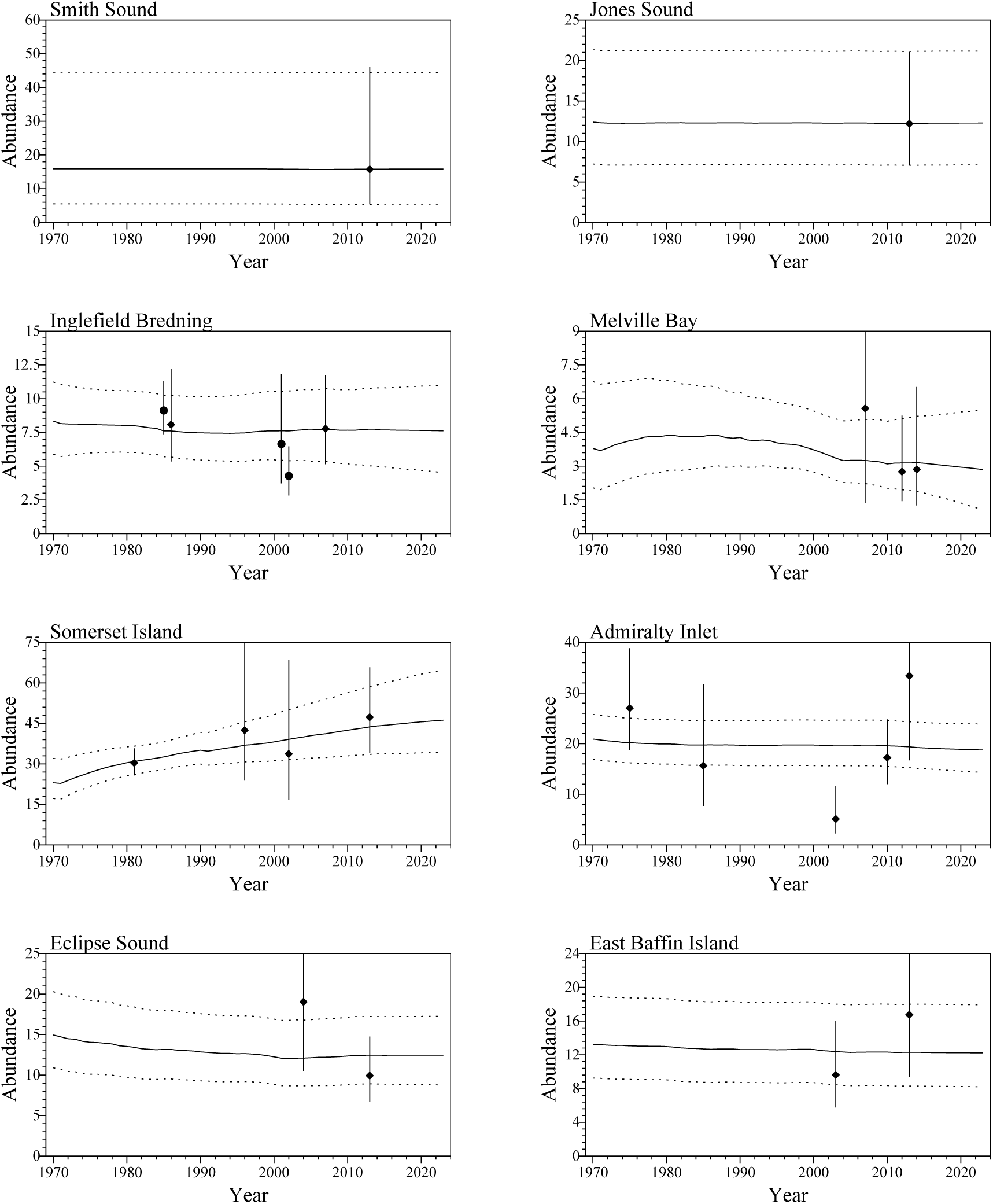
The trajectories of the different narwhal aggregations. Points with bars are the abundance estimates with 90% CI, solid curves the median, and dotted curves the 90% CI, of the estimated models. Abundance is given in thousands.

**Table 6:** Parameter estimates for the different models (*M*). Estimates are given by the median (*x*.5) and the 90% credibility interval (*x*.05 - *x*.95) of the posterior distributions. Abundance is given in thousands.

| $M$ | | $N_0$ | $N^*$ | $r$ | msyr | $p$ | $p_0$ | $a_m$ | $\gamma$ | msyl | $c_h$ |
| --- | --- | --- | --- | --- | --- | --- | --- | --- | --- | --- | --- |
| smith | $x_{.5}$ | - | 16 | .035 | .028 | .97 | .75 | 11 | 3 | .67 | .05 |
| | $x_{.05}$ | - | 5.5 | .013 | .01 | .96 | .53 | 8.1 | 2.1 | .62 | .0062 |
| | $x_{.95}$ | - | 45 | .058 | .045 | .99 | .97 | 14 | 3.9 | .7 | .29 |
| jones | $x_{.5}$ | - | 12 | .035 | .027 | .97 | .75 | 11 | 3 | .67 | .29 |
| | $x_{.05}$ | - | 7.2 | .013 | .0097 | .96 | .52 | 8.1 | 2.1 | .62 | .17 |
| | $x_{.95}$ | - | 21 | .058 | .046 | .99 | .97 | 14 | 3.9 | .7 | .6 |
| ingle | $x_{.5}$ | 8.3 | 11 | .023 | .018 | .97 | .66 | 12 | 2.9 | .65 | .12 |
| | $x_{.05}$ | 5.9 | 7.9 | .0055 | .0042 | .95 | .51 | 8.3 | 2.1 | .6 | .033 |
| | $x_{.95}$ | 11 | 25 | .056 | .044 | .99 | .95 | 14 | 3.9 | .7 | .48 |
| melvi | $x_{.5}$ | 3.8 | 7.6 | .037 | .029 | .97 | .77 | 11 | 3 | .67 | .16 |
| | $x_{.05}$ | 2 | 4.4 | .015 | .012 | .96 | .53 | 8 | 2.1 | .62 | .026 |
| | $x_{.95}$ | 6.8 | 25 | .06 | .047 | .99 | .98 | 14 | 3.9 | .71 | .49 |
| somer | $x_{.5}$ | 23 | 51 | .038 | .029 | .97 | .77 | 11 | 3 | .67 | .36 |
| | $x_{.05}$ | 17 | 36 | .022 | .017 | .96 | .54 | 8 | 2.1 | .62 | .075 |
| | $x_{.95}$ | 32 | 83 | .058 | .045 | .99 | .98 | 14 | 3.9 | .71 | .82 |
| admir | $x_{.5}$ | - | 21 | .035 | .027 | .97 | .74 | 11 | 3 | .66 | .37 |
| | $x_{.05}$ | - | 17 | .012 | .0092 | .96 | .52 | 8 | 2.1 | .61 | .086 |
| | $x_{.95}$ | - | 26 | .057 | .045 | .99 | .97 | 14 | 3.9 | .7 | .7 |
| eclip | $x_{.5}$ | - | 15 | .036 | .028 | .97 | .76 | 11 | 3 | .67 | .38 |
| | $x_{.05}$ | - | 11 | .013 | .01 | .96 | .53 | 8 | 2.1 | .62 | .077 |
| | $x_{.95}$ | - | 20 | .06 | .046 | .99 | .98 | 14 | 3.9 | .71 | .75 |
| baffi | $x_{.5}$ | - | 13 | .036 | .028 | .97 | .76 | 11 | 3 | .67 | .52 |
| | $x_{.05}$ | - | 9.2 | .015 | .011 | .96 | .53 | 8 | 2.1 | .62 | .2 |
| | $x_{.95}$ | - | 19 | .058 | .045 | .99 | .98 | 14 | 3.9 | .7 | .77 |

| $M$ | $N_t$ | $d_t$ | $\dot{r}_t$ | $\beta_a$ |
| --- | --- | --- | --- | --- |
| smith | 16 | 1 | .00029 | - |
|  | 5.4 | .98 | 6e-5 | - |
|  | 44 | 1 | .0011 | - |
| jones | 12 | .99 | .0011 | - |
|  | 7.1 | .97 | .00058 | - |
|  | 21 | 1 | .0019 | - |
| ingle | 7.7 | .7 | .012 | .32 |
|  | 4.8 | .25 | .0047 | .23 |
|  | 11 | .95 | .02 | .46 |
| melvi | 3.1 | .39 | .032 | - |
|  | 1.6 | .11 | .014 | - |
|  | 5.3 | .78 | .049 | - |
| somer | 45 | .92 | .0096 | - |
|  | 34 | .59 | .0052 | - |
|  | 61 | .98 | .022 | - |
| admir | 19 | .92 | .0089 | - |
|  | 15 | .8 | .0051 | - |
|  | 24 | .96 | .013 | - |
| eclip | 12 | .87 | .013 | - |
|  | 8.9 | .63 | .0079 | - |
|  | 17 | .96 | .02 | - |
| baffi | 12 | .94 | .0076 | - |
|  | 8.3 | .83 | .0046 | - |
|  | 18 | .98 | .012 | - |

**Figure 4:**
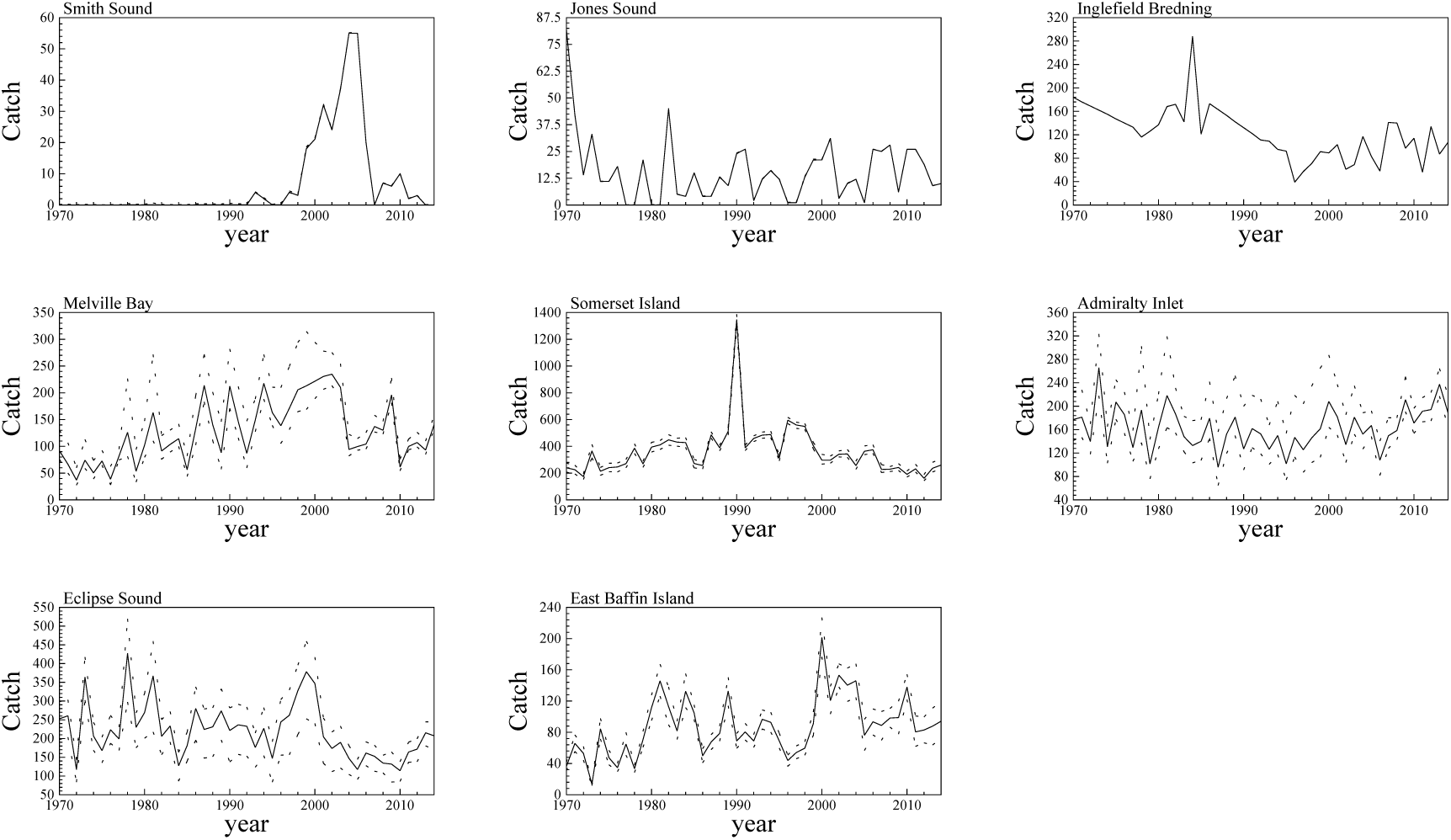
Yearly catches per summer aggregation with 90% confidence intervals.

Table 7 list the estimated total allowable takes for the different summer aggregations that will meet the management objective of Eq. (12) with probabilities from 0.5 to 0.95.

But management should define the total allowable takes for the different hunts (region and season), as these cannot generally be allocated directly to the different summer aggregation. Hence, Table 8 define possible total allowable takes for the different hunts, with Table 9 giving the associated estimates of the probabilities that these takes from 2015 to 2020 will allow the management objective to be fulfilled for the different summer aggregations. These latter probability estimates have 90% confidence limits that reflect the uncertainty of the summer aggregation origin of the animals taken in the different hunts. The C0 option in Table 8 is the average take over the five-year period from 2009 to 2013.

**Table 7:** Catch objective trade-off per stock. The total annual removals per stock that meet given probabilities (*P*) of meeting management objectives. The simulated period is from 2017 to 2022, and *F* is the assumed fraction of females in the catch.

| $P$ | smith | jones | ingle | melvi | somer | admir | eclip | baffi |
| --- | --- | --- | --- | --- | --- | --- | --- | --- |
| $F$ | 0.50 | 0.50 | 0.50 | 0.50 | 0.50 | 0.50 | 0.50 | 0.50 |
| 0.50 | 250 | 196 | 117 | 110 | 930 | 328 | 241 | 218 |
| 0.55 | 225 | 182 | 109 | 103 | 896 | 308 | 230 | 206 |
| 0.60 | 200 | 169 | 101 | 96 | 863 | 288 | 217 | 194 |
| 0.65 | 180 | 156 | 91 | 88 | 828 | 268 | 206 | 181 |
| 0.70 | 159 | 144 | 81 | 79 | 791 | 247 | 194 | 170 |
| 0.75 | 141 | 130 | 71 | 70 | 754 | 226 | 181 | 158 |
| 0.80 | 121 | 116 | 60 | 60 | 714 | 206 | 168 | 144 |
| 0.85 | 101 | 101 | 47 | 49 | 671 | 181 | 150 | 127 |
| 0.90 | 80 | 84 | 29 | 37 | 614 | 152 | 129 | 109 |
| 0.95 | 57 | 61 | 13 | 20 | 538 | 111 | 101 | 83 |

**Table 8:** Catch option examples (C#) of maximum yearly removal per hunting region.

| Hunt | Season | C0 | C1 |
| --- | --- | --- | --- |
| Etah | Spring | 4 | 5 |
| Qaanaaq | Summer | 98 | 98 |
| Grise Fjord | Spring | 7 | 9 |
| Grise Fjord | Summer | 11 | 15 |
| Grise Fjord | Fall | 0 | 0 |
| Upernavik | Summer | 100 | 70 |
| Ummannaq | Fall | 86 | 154 |
| Disko Bay | Winter | 73 | 97 |
| CCA | Spring | 4 | 6 |
| CCA | Summer | 74 | 118 |
| CCA | Fall | 2 | 3 |
| Arctic Bay | Spring | 31 | 41 |
| Arctic Bay | Summer | 141 | 188 |
| Arctic Bay | Fall | 0 | 0 |
| Pond Inlet | Spring | 58 | 77 |
| Pond Inlet | Summer | 55 | 73 |
| Pond Inlet | Fall | 4 | 5 |
| BIC | Spring | 12 | 11 |
| BIC | Summer | 100 | 91 |
| BIC | Fall | 44 | 40 |
| BIS | Spring | 5 | 5 |
| BIS | Summer | 9 | 8 |
| BIS | Fall | 12 | 11 |
| BIS | Winter | 0 | 0 |

**Table 9:** Examples of future annual removals (C#) per summer aggregation, with associated probabilities (P#) of fulfilling management objectives. The different removals follow from the catch options in Table 8, and the 90% confidence intervals of the estimates are given by the sub and super scripts.

|  | Smith | Jones | Inglefield | Melville | Somerset | Admiralty | Eclipse | Baffin |
| --- | --- | --- | --- | --- | --- | --- | --- | --- |
| C0 | 4 $\frac{4}{4}$ | 18 $\frac{18}{18}$ | 98 $\frac{98}{98}$ | 110 $\frac{133}{102}$ | 223 $\frac{246}{200}$ | 213 $\frac{260}{180}$ | 113 $\frac{141}{84}$ | 146 $\frac{158}{133}$ |
| P0 | 1.00 $\frac{1.00}{1.00}$ | 1.00 $\frac{1.00}{1.00}$ | 0.62 $\frac{0.62}{0.62}$ | 0.4 $\frac{0.45}{0.24}$ | 1.00 $\frac{1.00}{1.00}$ | 0.78 $\frac{0.85}{0.67}$ | 0.93 $\frac{0.96}{0.88}$ | 0.79 $\frac{0.83}{0.75}$ |
| C1 | 5 $\frac{5}{5}$ | 24 $\frac{24}{24}$ | 98 $\frac{98}{98}$ | 84 $\frac{114}{73}$ | 349 $\frac{376}{320}$ | 280 $\frac{338}{236}$ | 147 $\frac{183}{109}$ | 133 $\frac{144}{121}$ |
| P1 | 1.00 $\frac{1.00}{1.00}$ | 0.99 $\frac{0.99}{0.99}$ | 0.62 $\frac{0.62}{0.62}$ | 0.57 $\frac{0.64}{0.37}$ | 1.00 $\frac{1.00}{0.99}$ | 0.62 $\frac{0.72}{0.47}$ | 0.86 $\frac{0.94}{0.74}$ | 0.84 $\frac{0.87}{0.8}$ |

---

**Figure 5:**
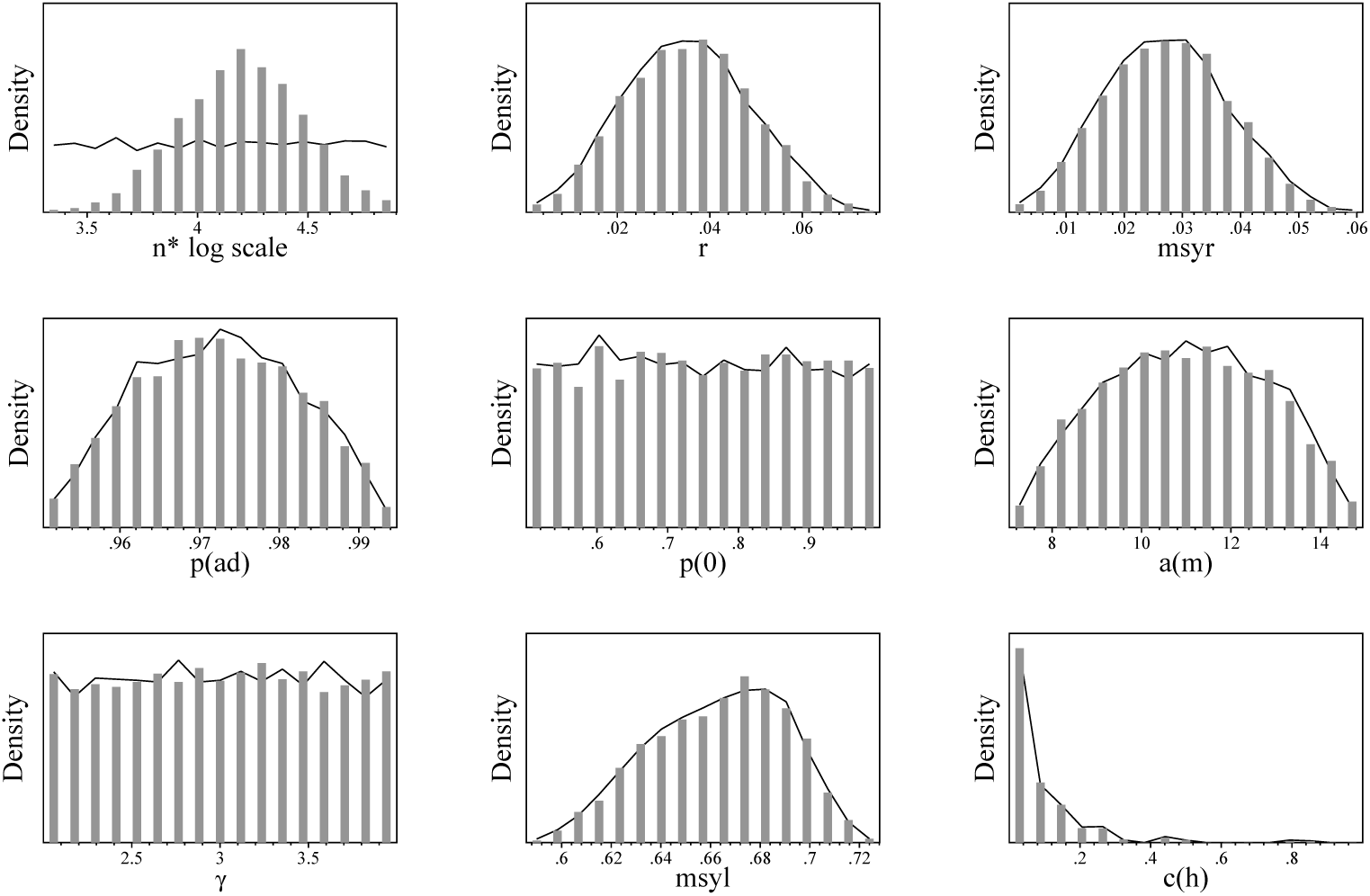
**Smith Sound** Realised prior (curve) and posterior (bars) distributions.

**Figure 6:**
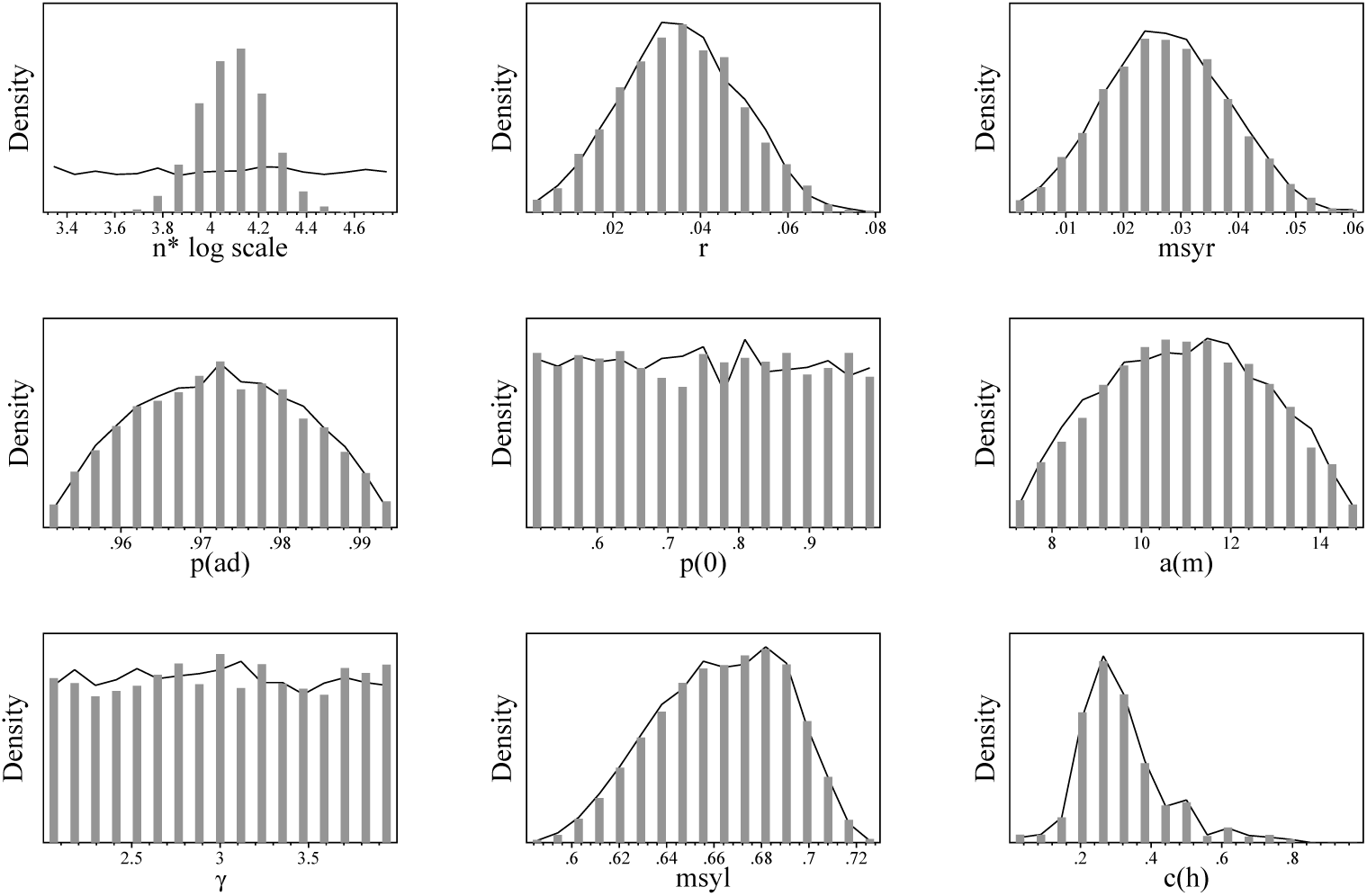
**Jones Sound** Realised prior (curve) and posterior (bars) distributions.

**Figure 7:**
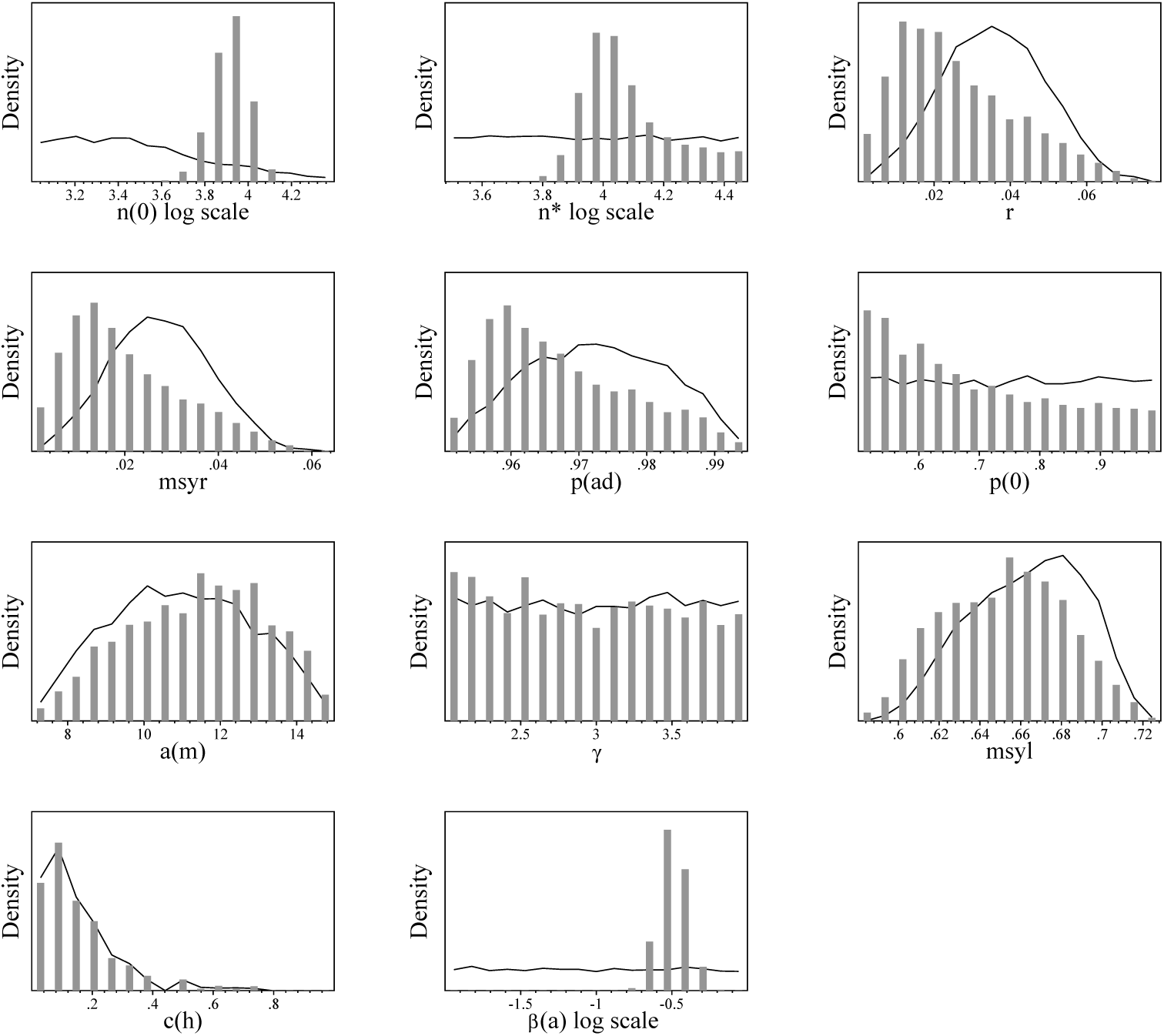
**Inglefield Bredning** Realised prior (curve) and posterior (bars) distributions.

**Figure 8:**
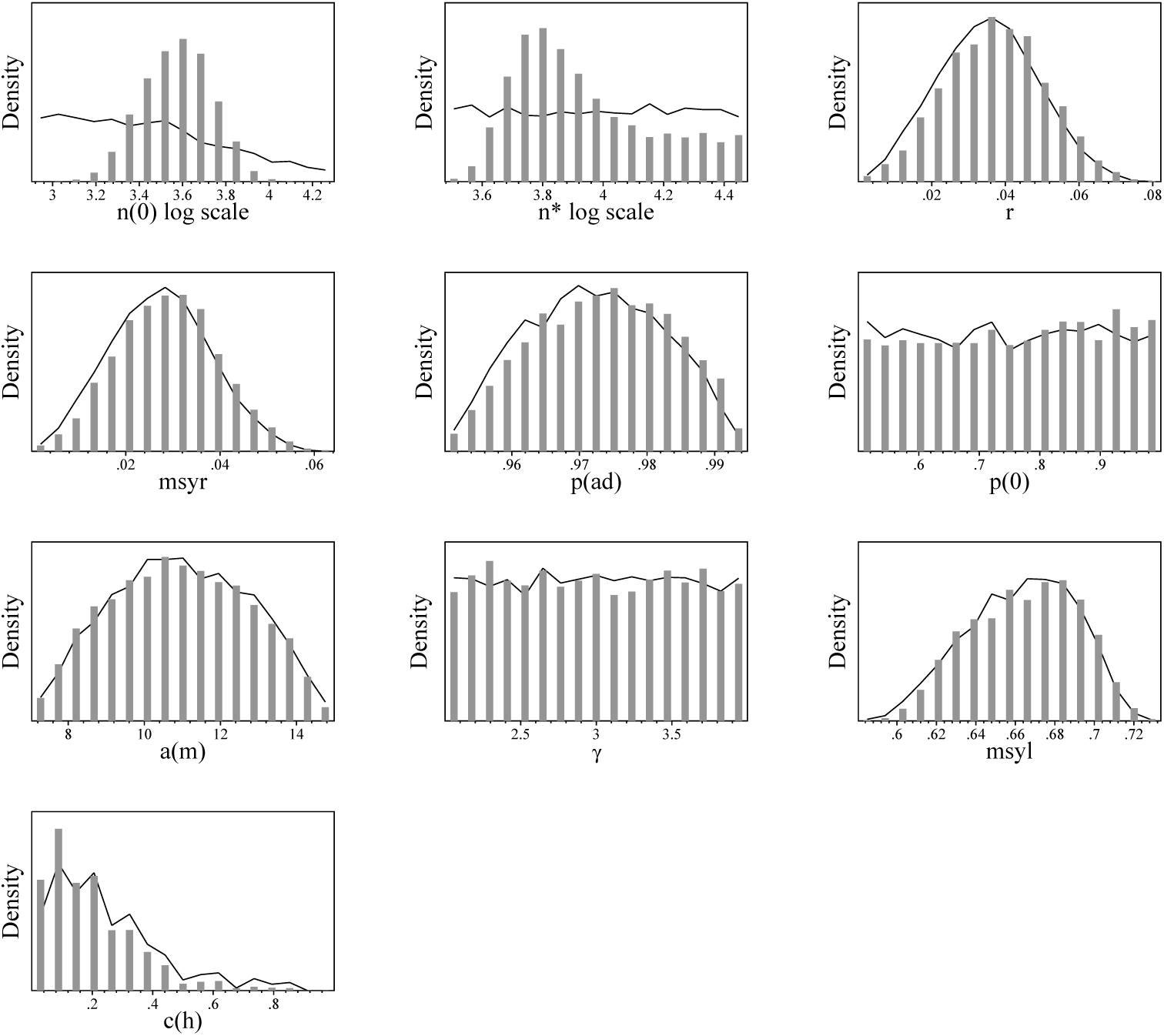
**Melville Bay** Realised prior (curve) and posterior (bars) distributions.

**Figure 9:**
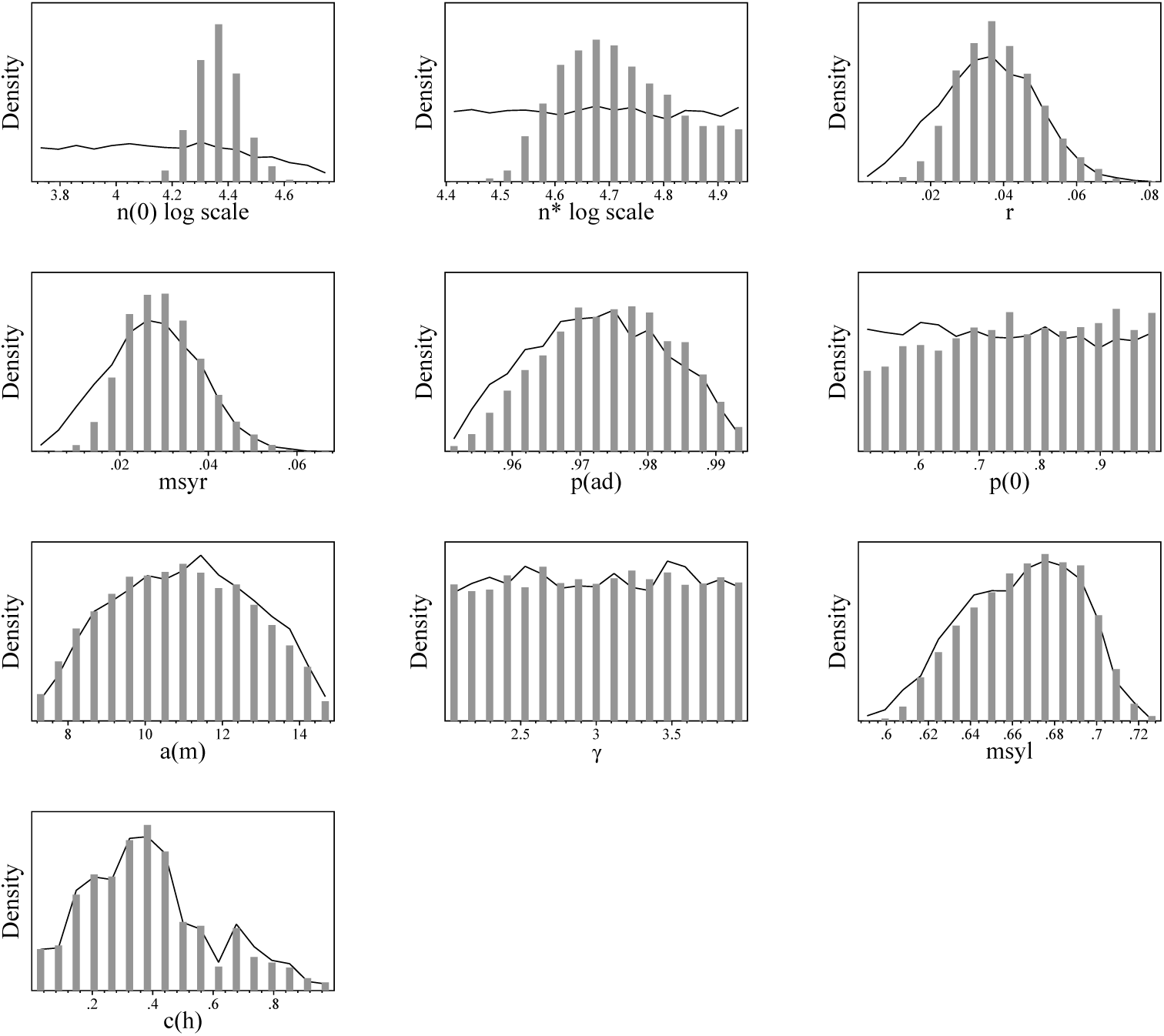
**Somerset Island** Realised prior (curve) and posterior (bars) distributions.

**Figure 10:**
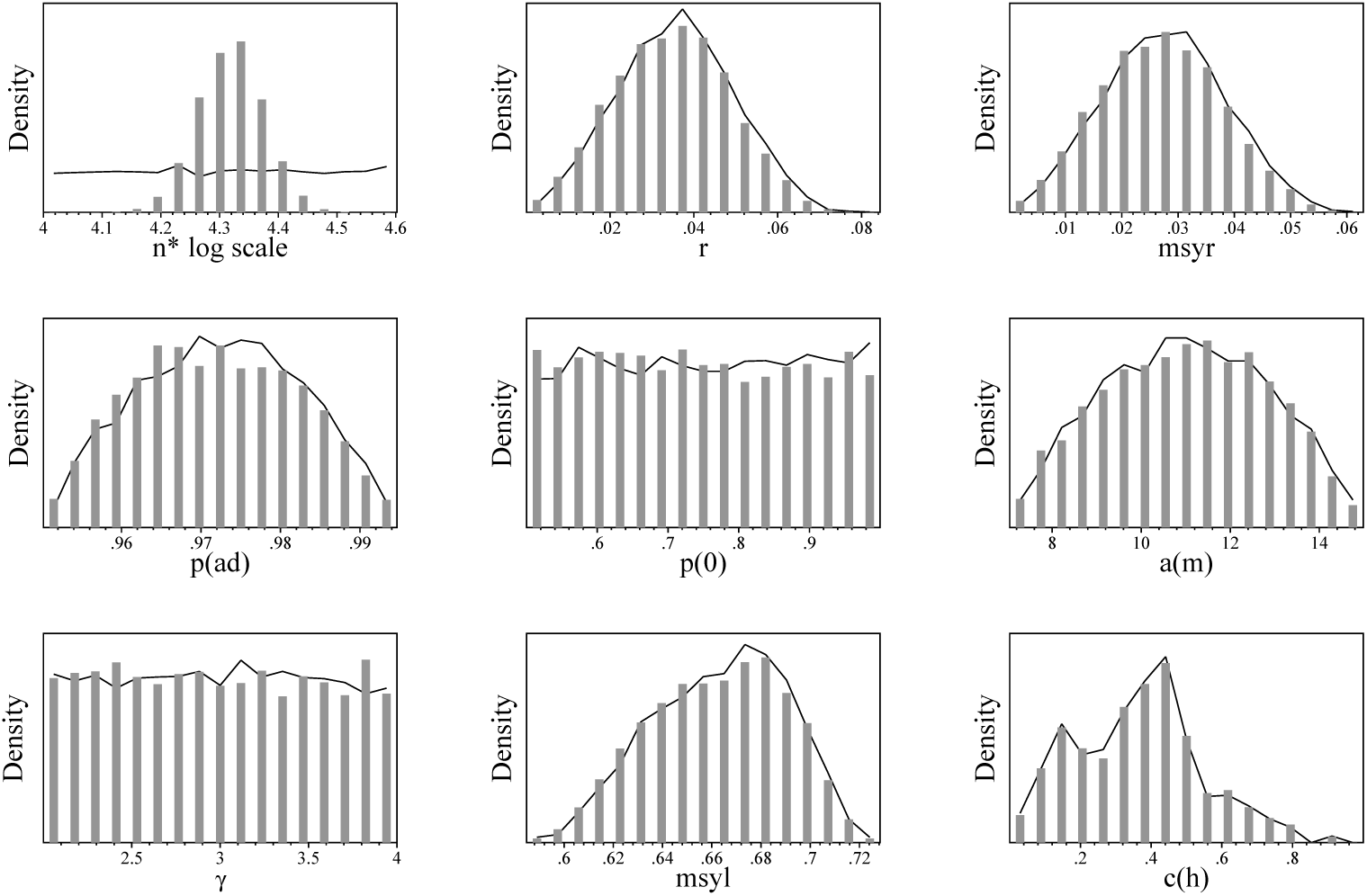
**Admiralty Inlet** Realised prior (curve) and posterior (bars) distributions.

**Figure 11:**
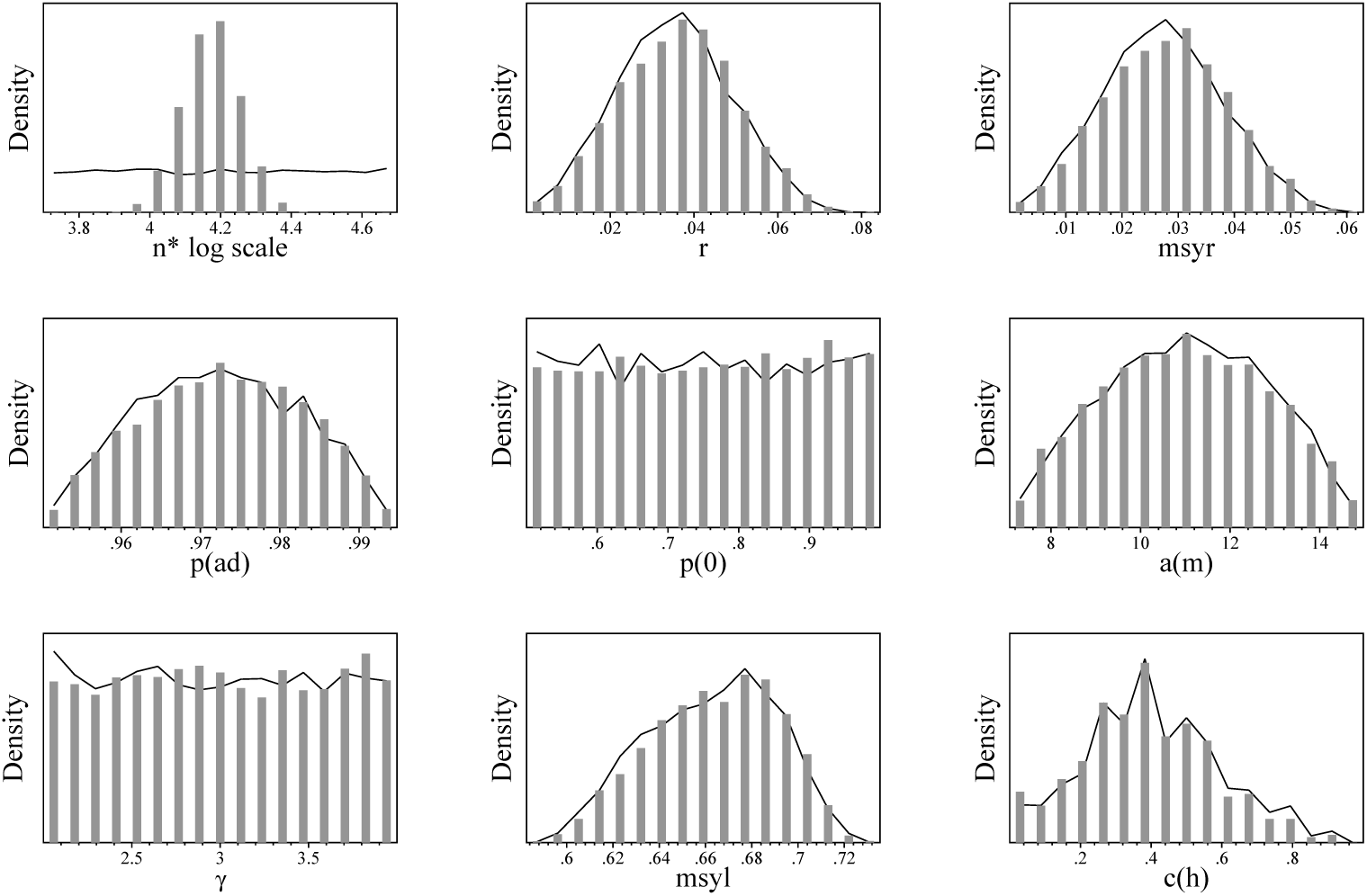
**Eclipse Sound** Realised prior (curve) and posterior (bars) distributions.

**Figure 12:**
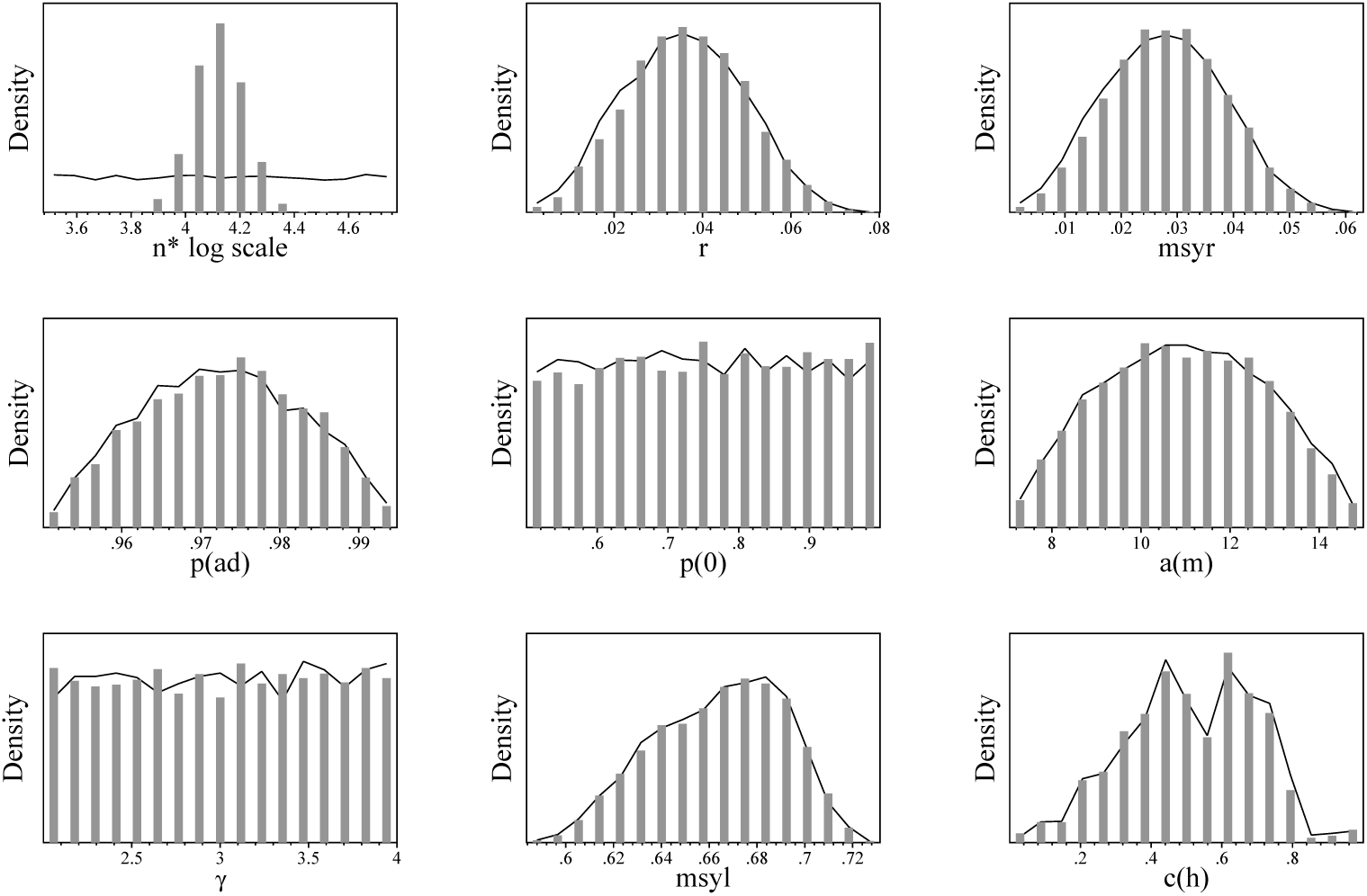
**East Baffin Island** Realised prior (curve) and posterior (bars) distributions.

**Table 10:**
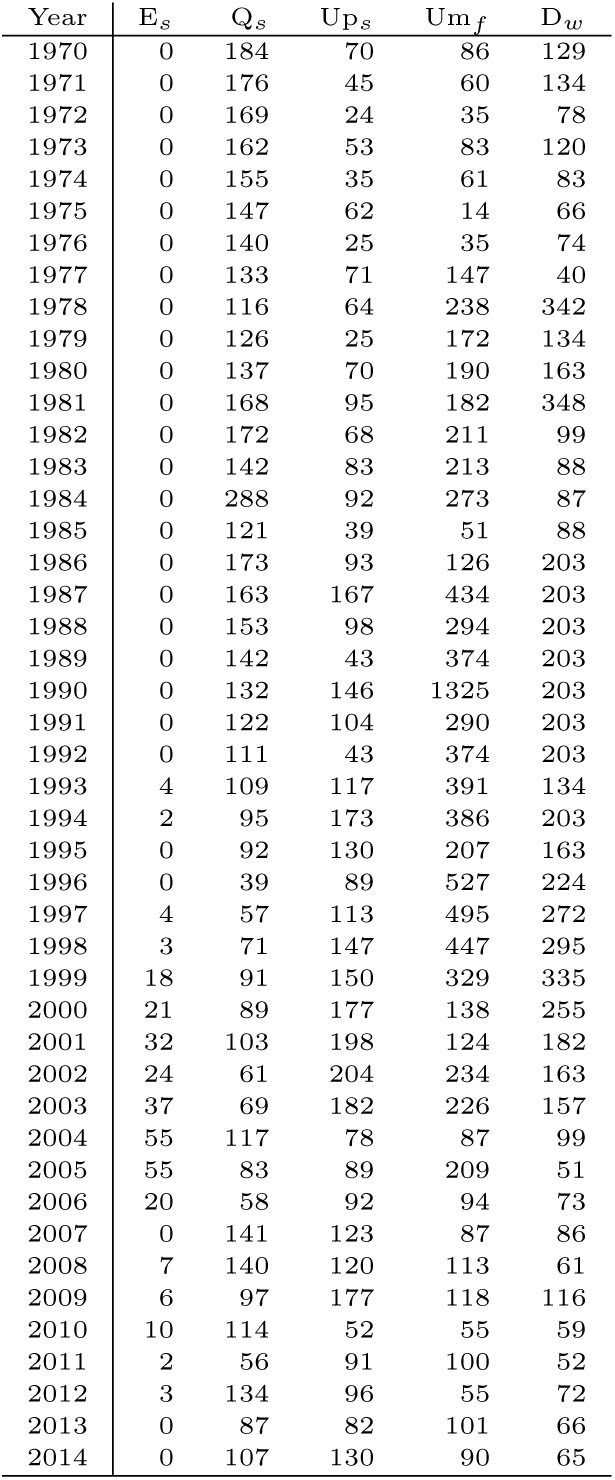
Estimated total removal per hunting region in Greenland per year. E*_s_*:Etah (Spring). Q*_s_*:Qaanaaq (Summer). Up*_s_*:Upernavik (Summer). Um*_f_*: Ummannaaq (Fall). D*_w_*:Disko Bay (Winter).

| Year | E<sub>s</sub> | Q<sub>s</sub> | Up<sub>s</sub> | Um<sub>f</sub> | D<sub>w</sub> |
| --- | --- | --- | --- | --- | --- |
| 1970 | 0 | 184 | 70 | 86 | 129 |
| 1971 | 0 | 176 | 45 | 60 | 134 |
| 1972 | 0 | 169 | 24 | 35 | 78 |
| 1973 | 0 | 162 | 53 | 83 | 120 |
| 1974 | 0 | 155 | 35 | 61 | 83 |
| 1975 | 0 | 147 | 62 | 14 | 66 |
| 1976 | 0 | 140 | 25 | 35 | 74 |
| 1977 | 0 | 133 | 71 | 147 | 40 |
| 1978 | 0 | 116 | 64 | 238 | 342 |
| 1979 | 0 | 126 | 25 | 172 | 134 |
| 1980 | 0 | 137 | 70 | 190 | 163 |
| 1981 | 0 | 168 | 95 | 182 | 348 |
| 1982 | 0 | 172 | 68 | 211 | 99 |
| 1983 | 0 | 142 | 83 | 213 | 88 |
| 1984 | 0 | 288 | 92 | 273 | 87 |
| 1985 | 0 | 121 | 39 | 51 | 88 |
| 1986 | 0 | 173 | 93 | 126 | 203 |
| 1987 | 0 | 163 | 167 | 434 | 203 |
| 1988 | 0 | 153 | 98 | 294 | 203 |
| 1989 | 0 | 142 | 43 | 374 | 203 |
| 1990 | 0 | 132 | 146 | 1325 | 203 |
| 1991 | 0 | 122 | 104 | 290 | 203 |
| 1992 | 0 | 111 | 43 | 374 | 203 |
| 1993 | 4 | 109 | 117 | 391 | 134 |
| 1994 | 2 | 95 | 173 | 386 | 203 |
| 1995 | 0 | 92 | 130 | 207 | 163 |
| 1996 | 0 | 39 | 89 | 527 | 224 |
| 1997 | 4 | 57 | 113 | 495 | 272 |
| 1998 | 3 | 71 | 147 | 447 | 295 |
| 1999 | 18 | 91 | 150 | 329 | 335 |
| 2000 | 21 | 89 | 177 | 138 | 255 |
| 2001 | 32 | 103 | 198 | 124 | 182 |
| 2002 | 24 | 61 | 204 | 234 | 163 |
| 2003 | 37 | 69 | 182 | 226 | 157 |
| 2004 | 55 | 117 | 78 | 87 | 99 |
| 2005 | 55 | 83 | 89 | 209 | 51 |
| 2006 | 20 | 58 | 92 | 94 | 73 |
| 2007 | 0 | 141 | 123 | 87 | 86 |
| 2008 | 7 | 140 | 120 | 113 | 61 |
| 2009 | 6 | 97 | 177 | 118 | 116 |
| 2010 | 10 | 114 | 52 | 55 | 59 |
| 2011 | 2 | 56 | 91 | 100 | 52 |
| 2012 | 3 | 134 | 96 | 55 | 72 |
| 2013 | 0 | 87 | 82 | 101 | 66 |
| 2014 | 0 | 107 | 130 | 90 | 65 |

**Table 11:** Estimated total removal per hunting region in Canada per year. G*_s_*:Grise Fjord (Spring). G*_s_*:Grise Fjord (Summer). G*_f_*: Grise Fjord (Fall). C*_s_*:CCA (Spring). C*_s_*:CCA (Summer). C*_f_*: CCA (Fall). A*_s_*:Arctic Bay (Spring). A*_s_*:Arctic Bay (Summer). A*_f_*:Arctic Bay (Fall). P*_s_*:Pond Inlet (Spring). P*_s_*:Pond Inlet (Summer). P*_f_*: Pond Inlet (Fall). B*_s_*:BIC (Spring). B*_s_*:BIC (Summer). B*_f_*:BIC (Fall). S*_s_*:BIS (Spring). S*_s_*:BIS (Summer). S*_f_*:BIS (Fall). S*_w_*:BIS (Winter).

| Year | G<sub>s</sub> | G<sub>s</sub> | G<sub>f</sub> | C<sub>s</sub> | C<sub>s</sub> | C<sub>f</sub> | A<sub>s</sub> | A<sub>s</sub> | A<sub>f</sub> | P<sub>s</sub> | P<sub>s</sub> | P<sub>f</sub> | B<sub>s</sub> | B<sub>s</sub> | B<sub>f</sub> | S<sub>s</sub> | S<sub>s</sub> | S<sub>f</sub> | S<sub>w</sub> |
| --- | --- | --- | --- | --- | --- | --- | --- | --- | --- | --- | --- | --- | --- | --- | --- | --- | --- | --- | --- |
| 1970 | 11 | 37 | 34 | 4 | 30 | 18 | 112 | 22 | 21 | 145 | 75 | 30 | 7 | 12 | 30 | 8 | 2 | 1 | 0 |
| 1971 | 6 | 19 | 17 | 4 | 32 | 21 | 112 | 23 | 21 | 141 | 74 | 30 | 10 | 20 | 43 | 26 | 4 | 3 | 0 |
| 1972 | 2 | 7 | 5 | 4 | 35 | 20 | 121 | 26 | 23 | 44 | 22 | 8 | 6 | 14 | 28 | 27 | 5 | 4 | 0 |
| 1973 | 3 | 18 | 12 | 4 | 62 | 30 | 183 | 37 | 33 | 264 | 136 | 60 | 3 | 6 | 11 | 0 | 1 | 0 | 0 |
| 1974 | 2 | 6 | 3 | 3 | 32 | 18 | 66 | 12 | 10 | 139 | 63 | 29 | 13 | 26 | 56 | 34 | 5 | 4 | 0 |
| 1975 | 2 | 6 | 3 | 6 | 36 | 19 | 220 | 35 | 30 | 107 | 46 | 25 | 4 | 12 | 17 | 34 | 4 | 4 | 0 |
| 1976 | 4 | 9 | 5 | 7 | 33 | 17 | 147 | 27 | 22 | 171 | 78 | 40 | 5 | 12 | 18 | 16 | 2 | 2 | 0 |
| 1977 | 0 | 0 | 0 | 0 | 13 | 36 | 23 | 22 | 19 | 144 | 69 | 34 | 0 | 14 | 111 | 6 | 0 | 0 | 0 |
| 1978 | 0 | 0 | 0 | 5 | 14 | 5 | 83 | 16 | 12 | 198 | 100 | 48 | 6 | 15 | 29 | 3 | 0 | 1 | 0 |
| 1979 | 0 | 2 | 19 | 0 | 4 | 15 | 42 | 6 | 8 | 118 | 97 | 0 | 25 | 6 | 23 | 23 | 27 | 3 | 0 |
| 1980 | 0 | 0 | 0 | 1 | 44 | 1 | 160 | 0 | 18 | 121 | 66 | 34 | 14 | 50 | 75 | 40 | 0 | 0 | 0 |
| 1981 | 0 | 0 | 0 | 0 | 20 | 86 | 113 | 34 | 20 | 101 | 58 | 29 | 15 | 52 | 77 | 53 | 8 | 23 | 0 |
| 1982 | 0 | 45 | 0 | 4 | 73 | 0 | 99 | 31 | 19 | 188 | 52 | 0 | 0 | 9 | 103 | 56 | 11 | 27 | 0 |
| 1983 | 0 | 3 | 2 | 0 | 64 | 29 | 141 | 18 | 14 | 81 | 71 | 78 | 38 | 42 | 36 | 4 | 2 | 0 | 0 |
| 1984 | 2 | 2 | 0 | 0 | 0 | 0 | 164 | 5 | 0 | 94 | 8 | 8 | 15 | 60 | 66 | 68 | 0 | 0 | 0 |
| 1985 | 7 | 6 | 2 | 11 | 18 | 8 | 183 | 0 | 0 | 141 | 27 | 62 | 8 | 78 | 5 | 36 | 2 | 0 | 0 |
| 1986 | 2 | 2 | 0 | 4 | 10 | 5 | 87 | 45 | 29 | 113 | 83 | 31 | 2 | 7 | 11 | 22 | 8 | 26 | 0 |
| 1987 | 2 | 2 | 0 | 4 | 11 | 6 | 22 | 11 | 7 | 57 | 44 | 17 | 9 | 35 | 65 | 0 | 0 | 0 | 0 |
| 1988 | 6 | 6 | 1 | 4 | 12 | 8 | 75 | 39 | 24 | 55 | 49 | 16 | 17 | 44 | 56 | 1 | 0 | 2 | 0 |
| 1989 | 4 | 4 | 1 | 7 | 21 | 11 | 86 | 46 | 27 | 74 | 74 | 25 | 15 | 47 | 74 | 28 | 9 | 34 | 0 |
| 1990 | 9 | 12 | 3 | 4 | 17 | 8 | 39 | 28 | 16 | 32 | 39 | 14 | 8 | 35 | 61 | 3 | 1 | 2 | 0 |
| 1991 | 10 | 13 | 3 | 4 | 18 | 15 | 65 | 49 | 26 | 46 | 55 | 21 | 10 | 40 | 66 | 3 | 2 | 5 | 0 |
| 1992 | 1 | 1 | 0 | 3 | 23 | 7 | 56 | 41 | 29 | 48 | 54 | 19 | 8 | 38 | 53 | 2 | 1 | 3 | 0 |
| 1993 | 5 | 6 | 1 | 5 | 30 | 10 | 46 | 36 | 22 | 38 | 43 | 15 | 9 | 43 | 65 | 12 | 5 | 15 | 0 |
| 1994 | 6 | 7 | 3 | 4 | 28 | 12 | 54 | 42 | 25 | 44 | 47 | 21 | 8 | 36 | 58 | 16 | 6 | 21 | 0 |
| 1995 | 5 | 5 | 2 | 3 | 26 | 12 | 33 | 9 | 16 | 90 | 0 | 0 | 8 | 34 | 61 | 3 | 1 | 4 | 0 |
| 1996 | 0 | 1 | 0 | 1 | 13 | 7 | 59 | 43 | 20 | 40 | 57 | 26 | 3 | 14 | 28 | 8 | 3 | 14 | 0 |
| 1997 | 0 | 1 | 0 | 2 | 21 | 12 | 40 | 29 | 13 | 28 | 43 | 21 | 5 | 26 | 57 | 1 | 0 | 1 | 0 |
| 1998 | 4 | 7 | 2 | 4 | 47 | 17 | 54 | 41 | 18 | 21 | 106 | 2 | 5 | 29 | 57 | 2 | 1 | 2 | 0 |
| 1999 | 8 | 11 | 2 | 3 | 12 | 8 | 49 | 45 | 16 | 17 | 106 | 39 | 1 | 29 | 79 | 14 | 4 | 25 | 0 |
| 2000 | 8 | 11 | 2 | 2 | 38 | 12 | 66 | 64 | 23 | 69 | 79 | 58 | 18 | 79 | 153 | 9 | 44 | 0 | 0 |
| 2001 | 12 | 16 | 3 | 6 | 54 | 37 | 67 | 71 | 24 | 27 | 32 | 21 | 13 | 53 | 108 | 11 | 1 | 14 | 0 |
| 2002 | 3 | 0 | 0 | 0 | 37 | 21 | 22 | 11 | 63 | 48 | 29 | 0 | 0 | 98 | 73 | 9 | 0 | 30 | 0 |
| 2003 | 0 | 10 | 0 | 4 | 32 | 2 | 60 | 84 | 15 | 32 | 40 | 10 | 12 | 73 | 105 | 36 | 0 | 1 | 0 |
| 2004 | 0 | 3 | 9 | 0 | 13 | 59 | 81 | 50 | 21 | 27 | 14 | 39 | 34 | 71 | 94 | 12 | 0 | 21 | 0 |
| 2005 | 1 | 0 | 0 | 0 | 43 | 26 | 83 | 93 | 1 | 26 | 25 | 20 | 14 | 10 | 133 | 0 | 0 | 6 | 0 |
| 2006 | 0 | 26 | 0 | 1 | 73 | 74 | 170 | 3 | 3 | 25 | 56 | 20 | 5 | 45 | 111 | 0 | 0 | 1 | 0 |
| 2007 | 4 | 21 | 0 | 0 | 44 | 12 | 90 | 72 | 5 | 8 | 35 | 32 | 10 | 31 | 120 | 4 | 1 | 0 | 0 |
| 2008 | 0 | 23 | 5 | 0 | 45 | 8 | 65 | 78 | 35 | 16 | 58 | 9 | 2 | 52 | 65 | 0 | 4 | 22 | 0 |
| 2009 | 5 | 1 | 0 | 3 | 22 | 46 | 23 | 150 | 1 | 24 | 6 | 21 | 9 | 25 | 93 | 10 | 0 | 40 | 0 |
| 2010 | 10 | 16 | 0 | 2 | 48 | 14 | 51 | 89 | 32 | 20 | 15 | 37 | 17 | 77 | 76 | 14 | 1 | 20 | 0 |
| 2011 | 14 | 10 | 2 | 8 | 51 | 4 | 38 | 112 | 26 | 45 | 81 | 3 | 7 | 23 | 125 | 2 | 0 | 4 | 0 |
| 2012 | 2 | 17 | 0 | 0 | 47 | 23 | 4 | 65 | 100 | 25 | 63 | 23 | 9 | 31 | 98 | 0 | 1 | 10 | 0 |
| 2013 | 5 | 0 | 4 | 1 | 33 | 23 | 43 | 167 | 4 | 30 | 82 | 58 | 11 | 9 | 143 | 3 | 1 | 18 | 0 |
| 2014 | 1 | 0 | 9 | 0 | 45 | 32 | 63 | 46 | 81 | 33 | 63 | 59 | 16 | 22 | 140 | 1 | 0 | 11 | 0 |

